# Asymmetric Redundancy of Soybean *Nodule Inception* (*NIN*) Genes in Root Nodule Symbiosis

**DOI:** 10.1101/2021.08.10.455861

**Authors:** Mengdi Fu, Jiafeng Sun, Xiaolin Li, Yuefeng Guan, Fang Xie

## Abstract

*NIN* is one of the most important root nodule symbiotic genes as it is required for both infection and nodule organogenesis in legume. Unlike most legumes with a sole *NIN* gene, there are four putative *NIN* genes in soybean (*Glycine max*). Whether and how these orthologs *NIN* genes contribute to soybean-rhizobia symbiotic interaction remain unknown. In this study, we found that all four *GmNIN* genes are induced by rhizobia, and that conserved CE and CYC binding motifs in their promoter regions are required for their expression in the nodule formation process. By generation of multiplex *Gmnin* mutants, we found that *Gmnin1a nin2a nin2b* triple mutant and *Gmnin1a nin1b nin2a nin2b* quadruple mutant displayed similar defects in rhizobia infection and root nodule formation, *Gmnin2a nin2b* produced less nodules but displayed hyper infection phenotype than wild type, while a *Gmnin1a nin1b* double mutant nodulated as wild type. Overexpression of *GmNIN1a*, *GmNIN1b, GmNIN2a*, and *GmNIN2b* reduced nodule numbers after rhizobia inoculation, with *GmNIN1b* overexpression having the weakest effect. In addition, overexpression of *GmNIN1a*, *GmNIN2a*, or *GmNIN2b*, but not *GmNIN1b*, produced malformed pseudo-nodule like structures without rhizobia inoculation. In conclusion, GmNIN1a, GmNIN2a and GmNIN2b play functionally redundant yet complicated roles for soybean nodulation. GmNIN1b, although is expressed at comparable level with other homologs, plays a minor role in root nodule symbiosis. Our work provides insight into the understanding of asymmetrically redundant function of *GmNIN* genes in soybean.

## INTRODUCTION

The legume-rhizobia symbiosis is the most important symbiotic association in terms of biological nitrogen fixation. In root nodules, endosymbiotic rhizobia use carbon from their legume hosts to fix atmospheric nitrogen into ammonia to provide the host plant with nitrogen nutrition. Understanding the underlying mechanisms of nodulation and nitrogen fixation is very important for improving agricultural production and for insights into natural ecosystems.

The legume-rhizobia symbiotic interaction is initiated by mutual recognition of molecular signals. The process is initiated by plant roots secreting flavonoid compounds into the soil, this attracts compatible rhizobia and stimulates them to synthesize and secrete highly specific lipochito-oligosaccharide (LCO) signalling molecules called nodulation factors (NFs) (Liu et al., 2016). Legume plants perceive NF signals via LysM receptor-like kinase expressed in their roots. NF recognition triggers two coordinated plant developmental programs: initiation of the infection process by which the bacteria enter the host cells, and, simultaneously, the elicitation of pericycle and cortical cell division resulting in nodule organ formation (Murray, 2011). When the bacteria reach the nodule primordium cells through the infection threads they are released into host-membrane enclosed structures called symbiosomes where they fix nitrogen (Oldroyd et al., 2011).

Soybean (*Glycine max*) is one of the most important legume crops in the world, and provides human food, animal feed, nutritional by-products and vegetable oil for human consumption and biofuel production. As a legume, soybean also has the capability to form root nodules through association with *Rhizobium* bacteria. The soybean genome was sequenced (Schmutz et al., 2010) and transcriptome and proteome analyses after rhizobia or NF inoculation have been performed in a number of studies (Hayashi et al., 2012; Libault et al., 2010; Libault et al., 2009; etc.). Understanding the biological function of the genes involved will help to establish elite cultivars that benefit sustainable farming practices. Forward and reverse genetic methods are available for soybean. However, due to multiple genome duplications that occurred at approximately 59 and 13 million years ago, approximately 75% of soybean genes are present in multiple copies leading to high genetic redundancy (Schmutz et al., 2010). Duplication occurred in segmental regions, accompanied by local genomic as well as sequence divergences. For example, the Nod factor receptor genes *NFR1* and *NFR5* are duplicated in soybean (GmNFR1α–GmNFR1β and GmNFR5α–GmNFR5β). GmNFR1α and GmNFR1β have 92% identity at the nucleotide level, but have different functions (Indrasumunar et al., 2010). On the other hand, GmNFR5α and GmNFR5β have 95% nucleotide identity and can functionally complement each other (Indrasumunar et al., 2011). The *Symbiosis Receptor Kinase* (*SymRK*) gene also has two copies in soybean (GmSymRKα and GmSymRKβ). RNAi knock down of the *GmSYMRK* genes revealed that *GmSymRKβ* is more important for root endosymbiosis than *GmSymRKα*, suggesting it retained its function after duplication (Indrasummunar et al., 2015).

*NIN* (*Nodule Inception*) is one of the most important genes in nodulation and is the founder gene for the NIN-like protein (NLP) transcription factor family in plants, which contain a conserved RWP-RK DNA-binding domain and a Phox and Bem1 (PB1) dimerization domain (Griesmann et al., 2018; Schauser et al., 1999; van Velzen et al., 2018). NIN has been shown to be required for rhizobial infection and nodule organogenesis in several legumes, including *M. truncatula*, *L. japonicus* and *Pisum sativum* (Marsh et al., 2007; Schauser et al., 1999; Borisov et al., 2003). LjNIN can directly bind to promoters of infection-specific genes and activate their expression, such as *NPL* (*Nodulation pectate lyase*), *SCARN* (*SCAR Nodulation*) and *EPR3* (*EPS receptor 3*) (Xie et al., 2012; Qiu et al., 2015; Kawaharada et al., 2017). NIN’s targets include genes which function in cortical cell division for root nodule organogenesis in *L. japonicus* and *M. truncatula*, including two *Nuclear Factor-Y* (*NF-Y*) subunit genes (*LjNF-YA1* and *LjNF-YB1*), the cytokinin receptor *MtCRE1*, and *LjLBD16a*/*ASL18* (Soyano et al., 2013; Vernie et al., 2015; Soyano et al., 2019). In addition, NIN has an essential role in the autoregulation of nodulation (AON) signaling pathway through its direct control of *CLE-Root Signal1/2* (*LjCLE-RS1/2*) or *MtCLE13/35* expression which act to limit nodule numbers (Soyano et al., 2014; Laffont et al., 2020; Luo et al., 2021). NIN can interact with other NLPs, including NLP1 which has important role in regulating nitrate inhibition of nodulation (Lin et al., 2018). Moreover, studies from non-legume nodulating plants, including *CgNIN* from the actinorhizal tree *Casuarina glauca* and *PaNIN* from *Parasponia*, showed that *NIN* is essential gene for actinorhizal and rhizobial nodulation, respectively (Clavijo et al., 2015; Bu et al., 2020). Thus, NIN is important for symbiotic nodulation, including rhizobial infection, nodule organogenesis, and the dynamic regulation of nodule numbers by nitrate through AON. In soybean, microarray data showed that several *NIN* genes were induced by rhizobia, but their function in nodulation has not been studied.

In this work, we investigated the expression patterns of the *GmNIN* genes, and used an RNAi knock downs strategy, CRISPR-Cas9 knock out mutants, and overexpression to fully analyze their function during the nodulation process. Our results show that GmNIN1a, GmNIN2a, and GmNIN2b function redundantly to mediate rhizobia infection and nodule initiation, and that GmNIN1b plays only a minor role.

## RESULTS

### *GmNINs* are Induced by Rhizobia and Have Nodulation-Specific Expression Patterns

The *G. max* genome encodes four GmNIN proteins belonging to the same clade as LjNIN and MtNIN (Lin et al., 2018) (Supplemental Figure S1). We then analyzed the gene expression patterns of the *GmNINs* by RT-qPCR after rhizobia inoculation. Our results showed that all four *GmNINs* can be induced by rhizobia and the induction levels were highest after 2 weeks post inoculation (Figure 1, A-D), and highly expressed in nodules (Figure 1, E-H). This result was consistent with previous microarray data that these *GmNINs* are highly induced during nodulation process (Libault et al.,2010).

**Figure 1.**
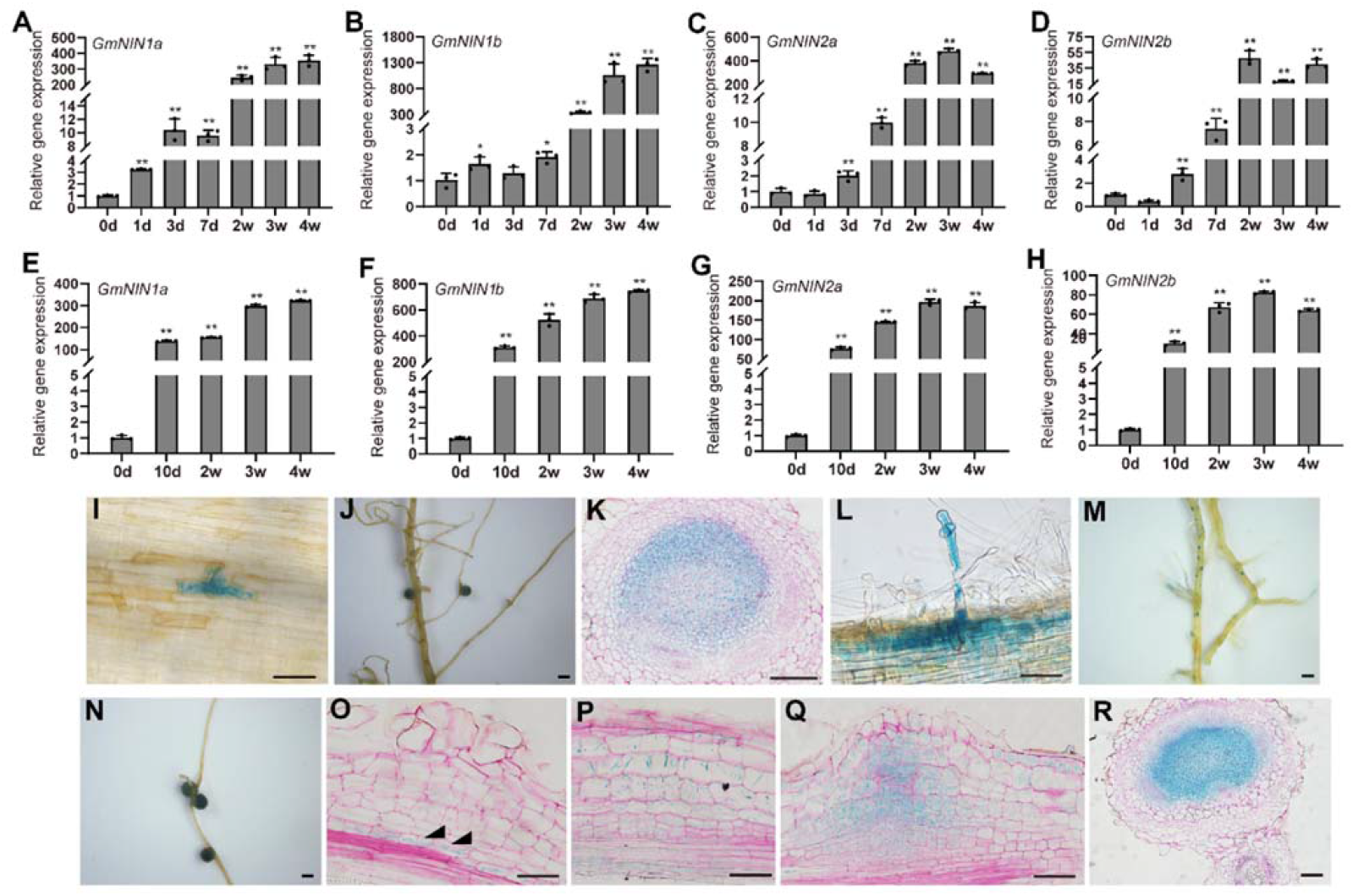
*GmNINs* were induced in soybean roots and nodules by rhizobia inoculation and have nodulation-specific expression patterns. Ato H, Expression levels of *GmNIN1a* (A, E), *GmNIN1b* (B, F), *GmNIN2a* (C, G) and *GmNIN2b* (D, H) in *G. max* roots (A-D) and nodules (E-H) after rhizobia inoculation. RT-qPCR were used to measure gene expression levels, normalized relative to uninoculated roots (0 d). Values are means ± SE. Data shown are from one representative experiment of three biological replicates. Asterisks indicate significant differences between treated and control conditions (0 d) (Student’s *t*-test, * *P*<0.05, ** *P*<0.01). I to R, The *pGmNIN2a_CYC_*:GUS (I-K), *pGmNIN2a_CE-CYC_*:GUS (L-R) were expressed in *G. max* hairy roots, and the images were taken at 3 days (I, L), 5 days (M) or 14 days (J, N) after rhizobia inoculation. (K, O-R) Section of these nodules stained with ruthenium red. Arrows (O) indicate GUS expression in pericycle cells. Scale bars: 1 mm (J, M and N); 100 μm (K, Q and R); 50 μm (I, L, O and P).

The promoter of *NIN* has a CYC motif (for CYCLOPS binding site) which is required for expression of *NIN* in epidermal cells to regulate rhizobial infection, and a remote conserved region with putative cytokinin response elements (CE region) which is required for expression of *NIN* in the pericycle to initiate nodule primordium formation in *L. japonicus* and *M. truncatula* (Singh et al., 2014; Liu et al., 2019).

Analysis of all four *GmNIN* promoter regions revealed CYC motifs within 3 kb upstream of their start codons (Supplemental Figure S2 and Table S1), and two (*GmNIN1a* and *GmNIN2a*) or three (*GmNIN1b* and *GmNIN2b*) conserved regions far upstream of their start codon (Supplemental Table S1). The 2^nd^ region was the most conserved region, which contains several putative cytokinin response elements in the promoter of *MtNIN* (Liu et al., 2019). Even all the 4 *GmNINs* contains the 2^nd^ CE region, it was conserved in *GmNIN1a, NIN2a and NIN2b,* but not in *GmNIN1b* promoter (Supplemental Figure S3). In order to further analyze the expression patterns of *GmNINs*, we made two promoter-GUS reporter constructs, one with the promoter of *GmNIN2a* containing its CYC motif (*pNIN2a_CYC_*) and a second version containing the 2^nd^ CE region and CYC motif (*pNIN2a_CE-CYC_*) fusion GUS reporter gene and expressed them in *G. max* by hairy root transformation. After rhizobia inoculation, weak GUS expression was detected in curled root hairs and nodules of *pNIN2a_CYC_:*GUS transgenic roots (Figure 1, I and J). In contrast, in roots transformed with the *pNIN2a_CE-CYC_*:GUS construct the GUS expression was much stronger, and staining was seen in root hairs, nodule primordia and nodules (Figure 1, L-N). Sectioning of the nodules revealed that both *pNIN2a_CYC_* and *pNIN2a_CE-CYC_* expressed in the nodule fixation zone, but the *pNIN2a_CE-CYC_* expression was stronger than *pNIN2a_CYC_* (Figure 1, K and R). However, only *pNIN2a_CE-CYC_*:GUS transgenic roots showed GUS activity in divided pericycle and cortex cells (Figure 1, O-Q). We further deleted (Δ) the D1, D2 or D3 in the 2^nd^ CE region, and found that ΔD1 displayed normal expression pattern, but only half of the transgenic plants can detect weaker GUS expression (Supplemental Figure S4, A-F and Table S2). However, ΔD2 or ΔD3 barely affected *GmNIN2a* expression pattern and expression level (Supplemental Figure S4, G-L, M-R and Table S2). This result was consistent with *MtNIN*, which showed that *pMtNIN*_CE(ΔD1)-CYC_:GUS cannot rescue *nin-1* nodulation phenotype (Liu et al., 2019), and suggested that the D1 motif is essential for *GmNINs* expression.

We then analyzed the GmNINs protein sequence, and found that the C-terminal RWP-RK and PB1 domain are conserved in the GmNINs, but the GmNIN1b N-terminal was shorter than other GmNINs (Supplemental Figure S5). Then the subcellular localization of GmNINs were examined using N-terminal GFP fusion proteins which were expressed in *N. benthamiana* leaves. Our result showed that these four GmNIN proteins displayed nuclear localization (Supplemental Figure S6).

### Silencing of *GmNINs* Using RNAi Reduced Nodule Numbers

In order to investigate the role of *GmNINs* in the root nodule symbiosis, we first used RNA interference (Ri) to reduce their transcript levels and nodule numbers were analyzed at 21 days post inoculation (dpi) with rhizobia. We found that *GmNIN1a Ri* plants produced no nodules; *GmNIN2a Ri* and *GmNIN2b Ri* transgenic roots had significantly reduced nodule numbers, while *GmNIN1b Ri* nodule numbers were less than the EV control but more than *GmNIN1a Ri*, *GmNIN2a Ri* and *GmNIN2b Ri* (Figure 2, A-F). The transcript levels of the *GmNINs* were analyzed by RT-qPCR, and the results showed that *GmNIN1a Ri* reduced expression of all four *GmNINs*, while *GmNIN1b Ri*, *GmNIN2a Ri* and *GmNIN2b Ri* reduced expression of just two of the four *GmNIN*s (Figure 2, G-H). We then analyzed *GmRIC1*, *GmRIC2* and *GmENOD40* genes expression levels in the *GmNINs Ri* roots. The RT-qPCR results showed that the expression of these genes was strongly reduced in the *GmNIN Ri* lines (Figure 2, I-J). These results suggest that these four *GmNINs* act redundantly in root nodule formation.

**Figure 2.**
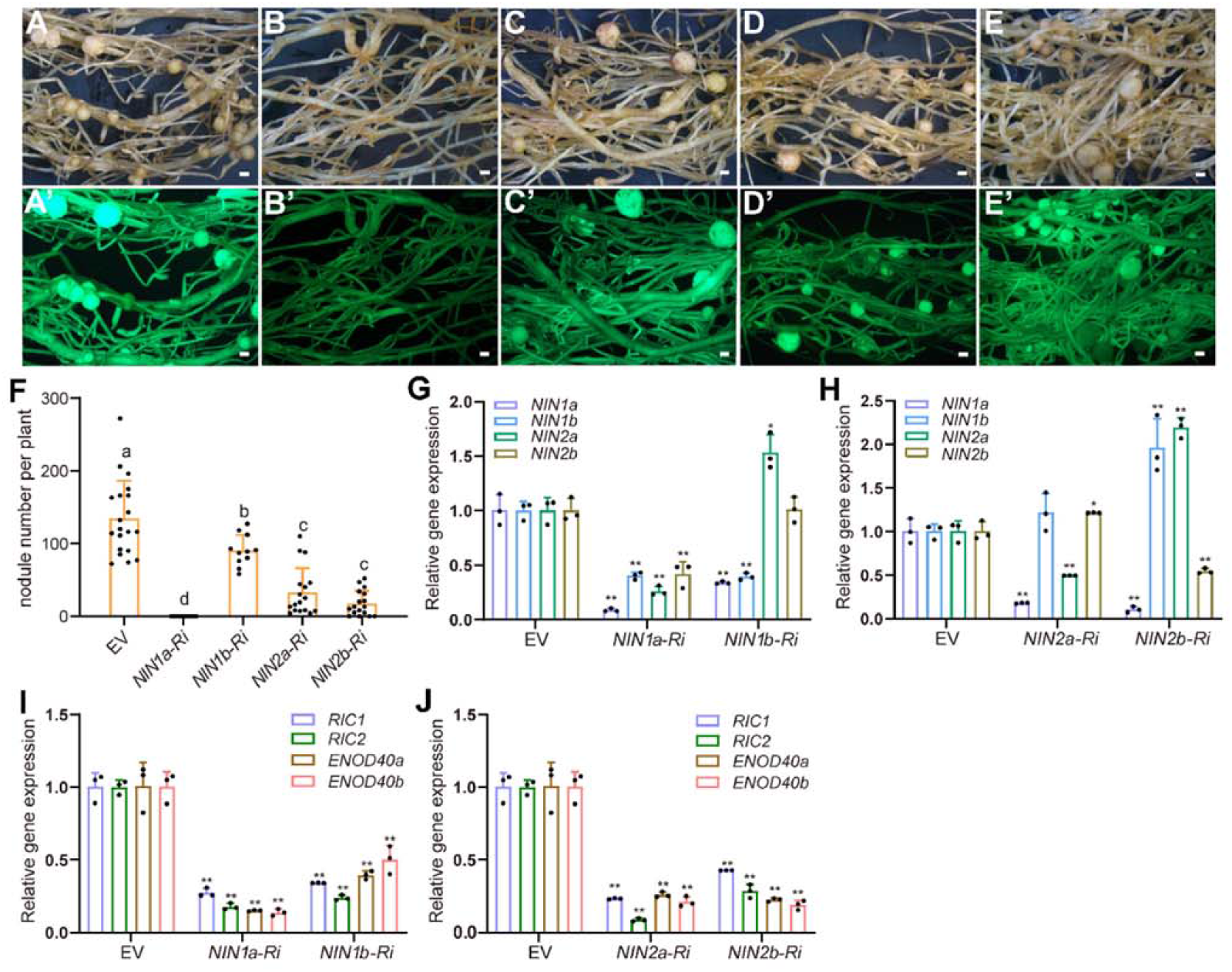
Nodulation phenotypes in *GmNIN* RNAi hairy roots. A to E, Nodule phenotypes of EV control (A), *GmNIN1a* (B), *GmNIN1b* (C), *GmNIN2a* (D) and *GmNIN2b* (E) RNAi transgenic roots 3 weeks after rhizobia inoculation. Upper panels (A-E) show bright-field images and the lower panels (A’-E’) are epifluorescence microscopy images showing GFP expression in the same transgenic roots. Scale bars = 1 mm. F, Nodule numbers of the EV and RNAi transgenic lines. The nodule numbers were scored at 3 weeks post inoculation with *B. japonicum* USDA110. Data from one representative experiment of three biological replicates. Error bars indicate SE, dots represent individual nodule numbers. The different letters indicate significant differences (one-way ANOVA-multiple comparisons). G to J, RT-qPCR analyze of the relative expression levels of *GmNINs* (G and H) and symbiotic reporter genes (I and J) in EV and *GmNINs* RNAi transgenic roots at 7 days post inoculation with *B. japonicum* USDA110. Error bars indicate SE. Data shown are from one representative experiment using three biological replicates. Asterisks indicate significant differences (\**P* < 0.05, \*\**P* < 0.01, Student’s *t*-test, comparisons between EV control and experimental groups).

### *GmNINs* CRISPR-Cas9 Mutants were Deficient in Rhizobia Infection and Nodule Organogenesis

To verify the function of *GmNINs* in the root nodule symbiosis process, we used multiplex mutagenesis via pooled CRISPR-Cas9 to generate high order *Gmnin* mutants (Bai et al., 2020). Vectors with small guide RNAs (sgRNAs) targeting the conserved RWP-RK motif or the N-terminus of *GmNIN1a/1b* or *GmNIN2a/2b* were constructed, respectively. These vectors were pooled transformed into soybean cultivar, Huachun 6, using *Agrobacterium tumefaciens*-mediated transformation. In T2 progenies, *Gmnin1b*, *Gmnin1a nin1b* and *Gmnin2a nin2b* double*, Gmnin1a nin2a nin2b* triple and *Gmnin1a nin1b nin2a nin2b* quadruple mutants were successfully obtained. These contained insertion or deletions that led to frameshifts which generated premature stop codons (Supplemental Figure S7).

The nodulation phenotypes of these mutants were further analyzed by inoculation with *B. japonicum* USDA110 which constitutively expresses the GUS reporter gene. *Gmnin1b* and *Gmnin1a nin1b* produced a similar number of nodules as wild type, *Gmnin2a nin2b* produced less nodule numbers than wild type, and *Gmnin1a nin2a nin2b* and *Gmnin1a nin1b nin2a nin2b* mutants did not develop any nodules at 21 dpi (Figure 3, A and B). RT-qPCR analyze revealed that *RIC1*, *RIC2* and *ENOD40* expression were reduced in these mutants, and the expression levels were much lower in the *Gmnin1a nin2a nin2b* and *Gmnin1a nin1b nin2a nin2b* mutants (Figure 3, C-E).

**Figure 3.**
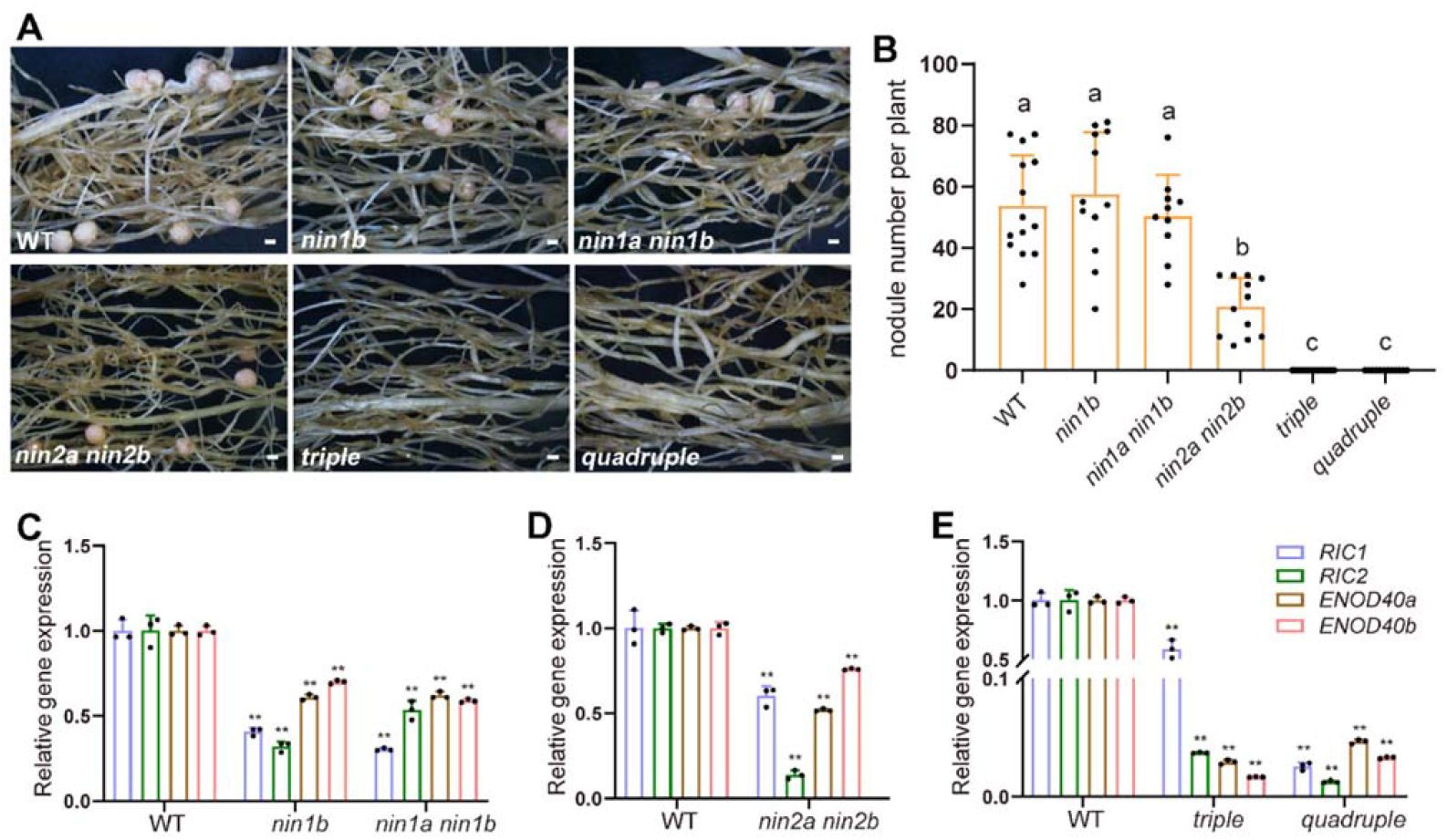
Nodulation phenotypes of *Gmnin* CRISPR-Cas9 mutants and gene expression levels in these mutants. A, Nodule phenotypes of *G. max* wild type (Huachun 6) and *Gmnin* CRISPR-Cas9 mutants 3 weeks after inoculation of *B. japonicum* USDA110. Sale bars= 1 mm. B, Nodule numbers of *G. max* wild type and *Gmnin* CRISPR-Cas9 mutants at 3 wpi. Error bars show SE, dots represent individual nodule numbers. Data from one representative experiment of two or three independent experiments. The different letters indicate significant differences (one-way ANOVA-multiple comparisons). C to E, RT-qPCR analyze of the relative expression levels of symbiotic reporter genes in wild type and *Gmnin* mutants. Error bars indicate SE. Data from one representative experiment of three biological replicates. Asterisks indicate significant differences (Student’s *t*-test, \**P* < 0.05, \*\**P* < 0.01, comparisons between WT control and experimental groups).

We further analyzed the root hair deformation and rhizobial infection thread formation phenotypes of these *Gmnin* mutants. In wild type and *Gmnin1a nin1b,* the root hairs were deformed and curled (Figure 4, A and B), and similar infection events including infection foci and infection threads can be produced 7 days after inoculation (Figure 4, E-H and M). *Gmnin1a nin2a nin2b* and *Gmnin1a nin1b nin2a nin2b* showed excessive root hair curling in response to rhizobia (Figure 4, C and D), no infection threads developed, and very rarely a few infection foci formed (Figure 4, K and L). Surprisingly, *Gmnin2a nin2b* showed hype infection phenotype, produce more infection foci and infection threads than wild type (Figure 4, I-J and N). Based on these results we concluded that the *Gmnin* triple or quadruple mutants have similar phenotypes as reported for *Ljnin* and *Mtnin* mutants, which showed excessive root hair deformation, developed occasional infection foci, and were unable to form nodules.

**Figure 4.**
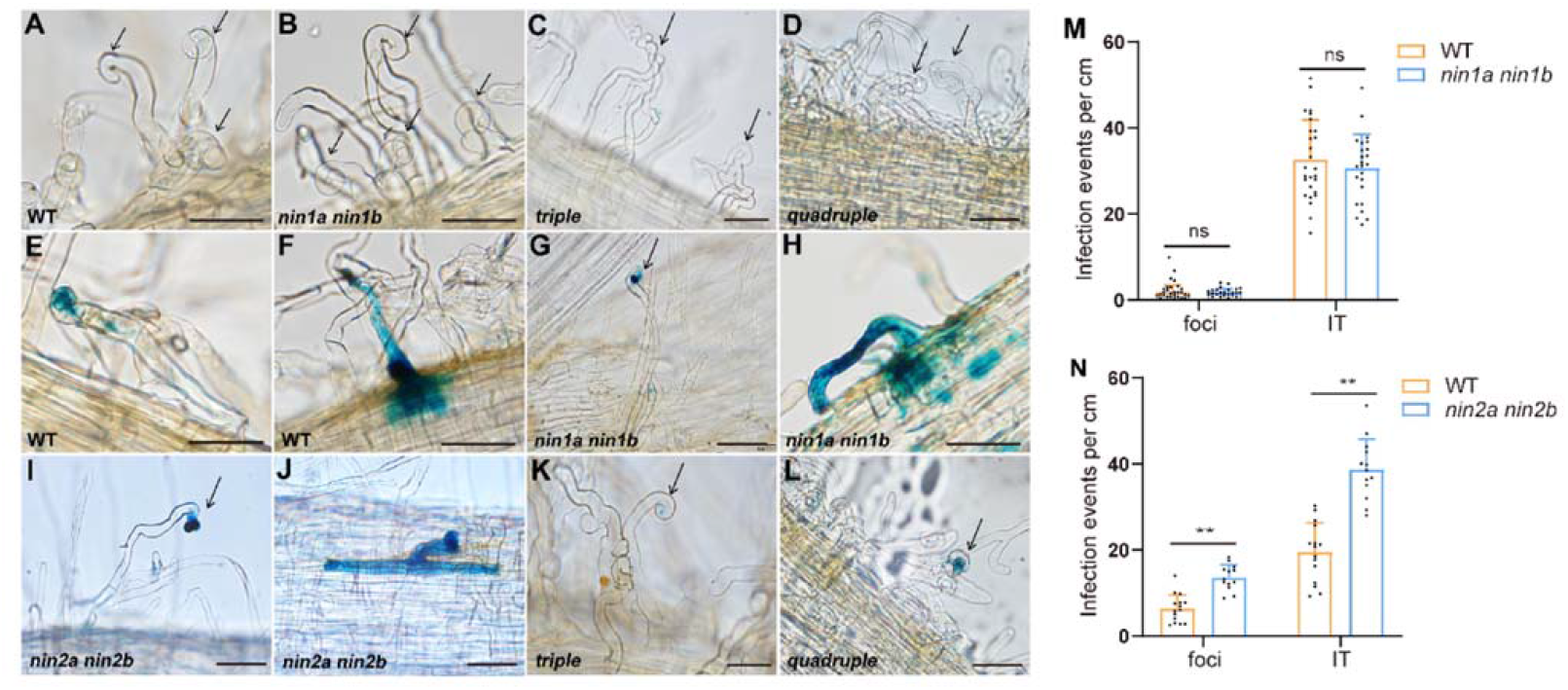
Root hair deformation and infection phenotypes of *Gmnin* mutants. A to D, Root hair deformation phenotypes of wild type (A), *Gmnin1a nin1b* (B), *Gmnin1a nin2a nin2b* (C) and *Gmnin1a nin1b nin2a nin2b* (D). E to L, Wild type (E, F), *Gmnin1a nin1b* (G, H) and *Gmnin2a nin2b* (I, J) plants have normal infection foci (E, G and I) and infection threads (F, H and J), but *Gmnin1a nin2a nin2b* (K) and *Gmnin1a nin1b nin2a nin2b* (L) only occasionally developed a few infection foci after rhizobia inoculation. Arrowheads represent deformed root hairs. Scale bars = 100 μm. M to N, Infection events in wild type and *Gmnin1a nin1b* (M)*, Gmnin2a nin2b* (N) 7 days after inoculation of *B. japonicum* USDA110 carrying a GUS reporter gene. Error bars show SE, dots represent individual numbers. Data from one representative experiment of two independent experiments. Foci: infection foci; IT: infection thread. ns: no significant difference (Student’s *t*-test).

### Overexpression of *GmNINs* Blocked Nodulation but Produced Spontaneous Nodule Primordium-Like Structures

It has been shown that overexpression of *NIN* can reduce nodule numbers after rhizobia inoculation (Soyano et al., 2014),16 and that it can result in formation of some root nodule-like structures in the absence of rhizobia inoculation (Soyano et al., 2013; Vernie et al., 2015). We then overexpressed *GmNINs* using 35S promoter driven *GmNIN* cDNA in *G. max* hairy roots. The transgenic roots were inoculated with rhizobia and the nodulation phenotypes were scored at 21 dpi with *B. japonicum*. We found that overexpression of *GmNIN1a*, *GmNIN2a* or *GmNIN2b* dramatically reduced nodule numbers; while with overexpression *GmNIN1b,* the nodule numbers were less than EV control but more than *GmNIN1a* and *GmNIN2a* overexpression lines (Figure 5, A-D). The expression of symbiotic reporter genes were examined in the *GmNIN1a* overexpression roots after rhizobia inoculation. The results showed that *GmNIN1a* was highly induced after rhizobia inoculation and in *GmNIN1a* overexpression hairy roots (Figure 5E). Symbiotic reporter genes such as *RIC1*, *RIC2, ENOD40* and *NF-YA1* were induced by rhizobia in the EV control and the *GmNIN1a* overexpression transgenic roots (Figure 5, F-K).

**Figure 5.**
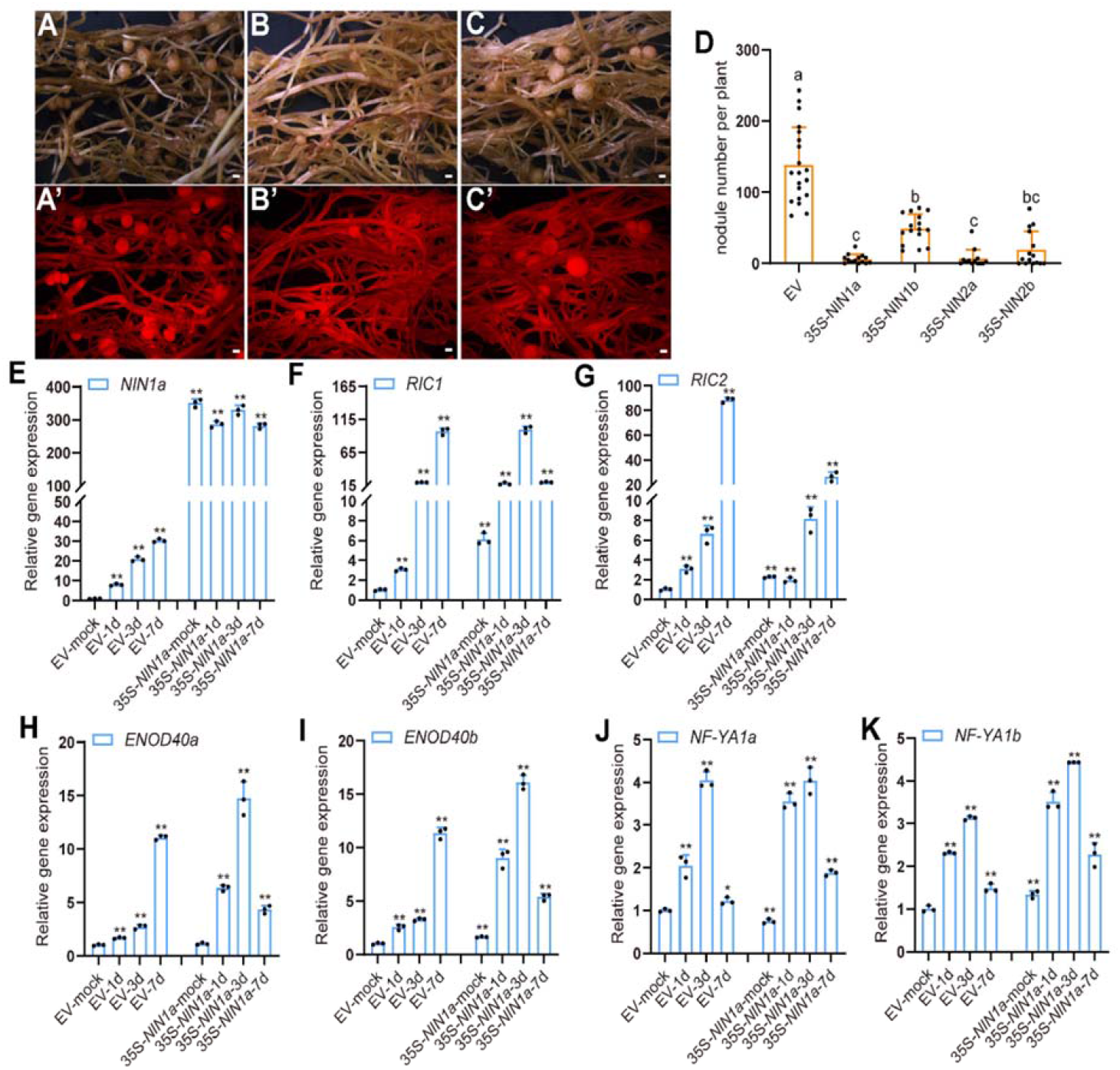
*GmNIN* overexpression lines suppress nodulation. A to C, Nodule phenotypes of EV (A), *p35S: GmNIN1a* (B) and *p35S: GmNIN1b* (C) lines at 3 wpi. Scale bars= 1 mm. Upper panels (A-C) represent bright-field images, and the lower panels (A’-C’) represent fluorescence images show mCherry expression in the same transgenic roots. D, Nodule numbers of *GmNIN* overexpression transgenic roots at 3 wpi with *B. japonicum*. Error bars show SE, dots represent individual numbers. Data from one representative experiment of three independent experiments. Different letters indicate significant differences (one-way ANOVA-multiple comparisons). E to K, RT-qPCR analyze transcription levels of *GmNIN1a* (E), *RIC1* (F), *RIC2* (G), *ENOD40a* (H), *ENOD40b* (I), *NF-YA1a* (J) and *NF-YA1b* (K) in EV and *GmNIN1a-OE* hairy roots after rhizobia inoculation. Error bars indicate SE. Data from one representative experiment of three biological replicates. Asterisks indicate significant differences (Student’s *t-*test, \**P* < 0.05, \*\**P* < 0.01, comparisons between EV control and *GmNIN1a-OE* hairy roots).

We next examined whether overexpression of *GmNINs* can induce the formation of nodule primordium-like structures without rhizobia inoculation. The composite plants with transgenic hairy roots were transferred intro vermiculite: perlite mixture (1:1) and the phenotype was scored 35 days after transfer. We found that overexpression of *GmNIN1a*, *GmNIN2a* and *GmNIN2b* produced abnormal root growths, including white nodule-like structures, small bumps and short but enlarged lateral roots (Figure 6, A-D, I and Supplemental Figure S8, C-H). Nearly all the *GmNIN1a* overexpression roots, 60% of *GmNIN2a* overexpression roots, and 80% of *GmNIN2b* overexpression roots showed these abnormal architectures, but none of EV and *GmNIN1b* overexpression roots developed this kind of malformed structures, in three independent experiments (Figure 6I and Supplemental Figure S8, A-H). Sectioning of the *GmNINs*-induced abnormal structures revealed that the nodule-like structures and white bumps were formed *via* cortical cell division and were anatomically similar to root nodule primordia (Figure 6, E-F and Supplemental Figure S8, I-J, M-N); the malformed nodule-like structures have central vascular systems similar to lateral roots but also have undergone extensive cortical cell divisions and show over-proliferation of pericycle and cortical cells (Figure 6, G-H and Supplemental Figure S8, K-L, O-P). RT-qPCR analysis showed that *GmNIN1a* was highly expressed in the transgenic hairy roots, but the early nodulin gene *ENOD40* was highly expressed in the roots showing abnormal architecture, while was not changed in normal roots (Figure 6, J-L).

**Figure 6.**
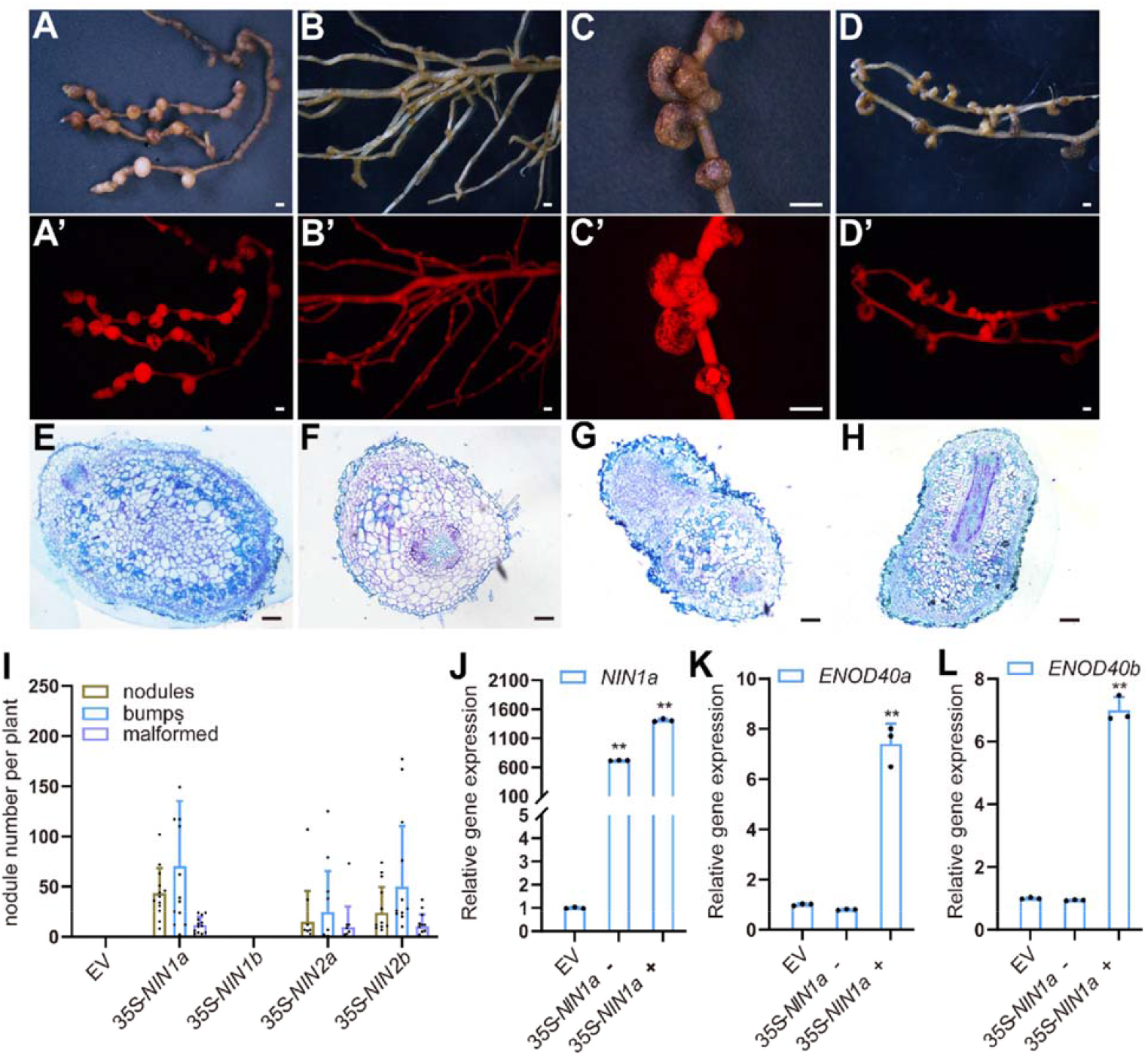
*GmNINs* overexpression triggers the formation of spontaneous nodule-like structures. A to D, *p35S: GmNIN1a* expressed in wild type hairy roots and cultured for 5 weeks without rhizobia inoculation, produced nodules (A), small white bumps (B) and malformed nodule-like structures (C and D). Upper panels (A-D) showed bright-field images, and lower panels (A’-D’) are epifluorescence microscopy images show mCherry expression in the same transgenic roots. Scale bars = 1 mm. E to H, Transverse sections of spontaneous nodules (E), small white bumps (F) and malformed nodule-like structures (G and H) in *GmNIN1a-OE* hairy roots in the absence of rhizobia inoculation. Scale bars = 100 μm. I, Nodule numbers in the overexpressed *GmNINs* hairy roots 5 weeks after culture. Dots represent individual numbers. Data from one representative experiment of three independent experiments. J to L, RT-qPCR analyze transcription levels of *GmNIN1a* (J), *ENOD40a* (K) and *ENOD40b* (L) in EV and *p35S:GmNIN1a* hairy roots. *p35S: NIN1a -* represent no spontaneous nodule-like structures hairy roots, and *p35S:* NIN1a + represent have spontaneous nodule-like structures hairy roots. Error bars indicate SE. Data from one representative experiment of three biological replicates. Asterisks indicate significant differences (Student’s *t*-test, \**P* < 0.05, \*\**P* < 0.01, comparisons between EV control and *GmNIN1a-OE* hairy roots).

LjNIN systemically suppresses nodulation through the AON pathway (Soyano et al., 2014). We expressed *35S:GmNIN1a* in wild type or *nark* mutant plants, and the nodulation phenotype was analyzed in transformed or untransformed hairy roots at 21 dpi. The mean number of nodules was significantly suppressed in both mCherry-positive transgenic and mCherry-negative untransformed roots in wild type background, but the repressive level was stronger in mCherry-positive transgenic roots (Figure 7, A-B and E). However, the suppression effect was not present in *nark* mutant background either in transgenic or non-transgenic hairy roots (Figure 7, C-D and E). This result indicates that ectopic expression of *GmNIN1a* systemically inhibits nodulation *via* the AON pathway.

**Figure 7.**
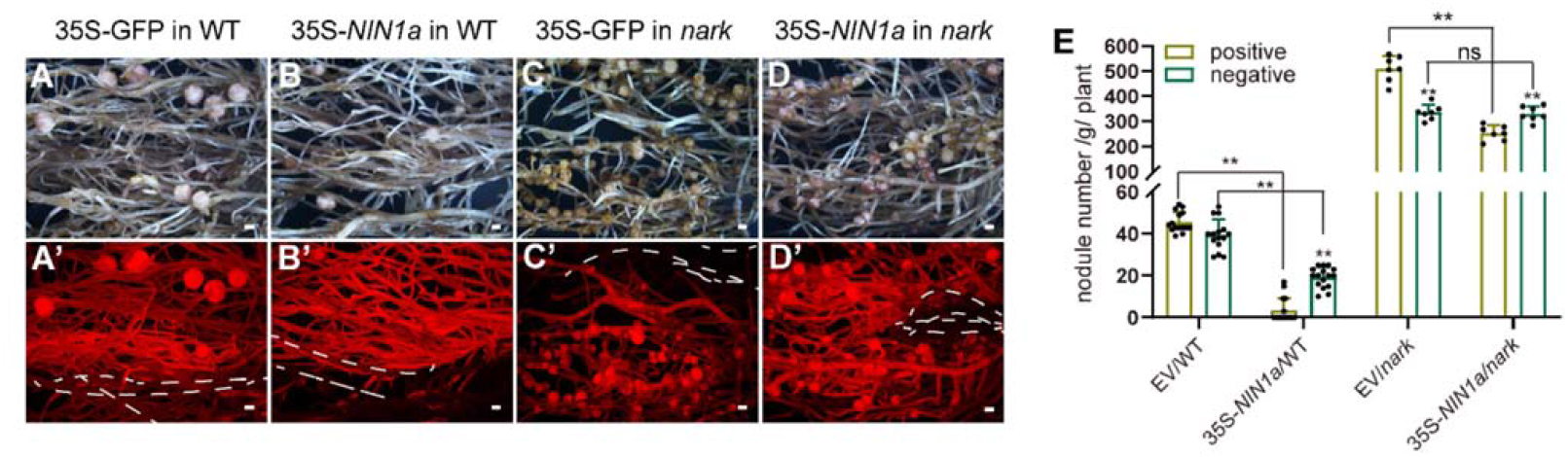
Systemic inhibition of nodulation by overexpression of *GmNIN1a*. Ato D, Nodulation on mCherry-negative untransformed roots and mCherry-positive transformed roots on wild type carrying EV (A) or *p35S-GmNIN1a* (B), and on *nark* carrying EV (C) or *p35S-GmNIN1a* (D) at 3 wpi. Upper panels (A-D) show composite bright-filed images. Lower panels (A’-D’) show corresponding fluorescent images with mCherry as a transformation marker for transgenic hairy roots. White dash in lower panels represent mCherry-negative roots. Scale bars= 1 mm. E, Nodule numbers on mCherry-positive and -negative roots that were transformed with the EV or *p35S-GmNIN1a* in wild type (WT) or *nark* mutants. Dots represent individual numbers. Asterisks indicate significant differences between mCherry-negative untransformed roots and mCherry-positive transformed roots on wild type carrying EV or *p35S-GmNIN1a* (Student’s *t*-test, \**P*<0.05, ** *P*<0.01, ns: no significant difference).

## DISCUSSION

Soybean is an ancient polyploid due to genome duplications, resulting in a nearly 75% of genes being present in multiple copies (Schmutz et al., 2010). This makes it more difficult to identify recessive inheritance in this duplicated genome. Using a CRISPR-Cas9 approach, we were able to make higher order mutants, and then analyzed their biological function. NIN is essential for all processes in root nodule formation, and *G. max* contains four *NIN* genes due to genome duplications, but which genes are required for nodulation was not studied. In this work, we carefully examined *GmNIN* expression patterns and nodulation phenotypes in CRISPR-Cas9 mutants or *GmNIN* overexpression lines. The results suggested that GmNIN1a, GmNIN2a and GmNIN2b are essential for nodulation in a functionally redundant manner, but that GmNIN1b likely has only a minor role in root nodule symbiosis.

*NIN* is specifically induced and expressed in nodules (Marsh et al., 2007; Schauser et al., 1999). In *G. max*, all these four *GmNINs* were induced by rhizobia and were highly expressed in nodules. In all these *GmNIN* promoters, contains the CYC motif, but the most important 2^nd^ CE motif was not conserved in *GmNIN1b* promoter. The CYC motif is required for *NIN’s* expression in epidermal cells where infection threads are initiated. Compared with p*MtNIN_CYC_*:GUS, which is expressed in epidermal and pericycle cells (Liu et al., 2019), p*GmNIN2aCYC*:GUS was only detected in epidermal cells and nodules. This difference might result from shorter promoter we used, and the GUS expression was weaker than detective. Or this might also suggests that another motif in *NIN* promoter besides the CYC motif may be required for *NIN* expression in pericycle cells. However, p*GmNIN2a_CE-CYC_*:GUS was found to express in epidermal cells, pericycle cells, divided cortical cells, and in nodules, suggesting that this conserved 2^nd^ CE motif is essential and sufficient for *GmNIN2a* expressed in pericycle cells.

Compared with other species, *G. max* contains four *NIN* genes and all of them were induced and expressed in nodules. Several studies including ours, showed that RNAi knock down *GmNIN1a* showed a nodule-minus phenotype (He et al., 2020 and this study), suggesting that *GmNIN1a* is sufficient for NIN’s function in root nodule symbiosis. However, after careful analysis of *GmNIN* expression levels in *GmNIN1a* RNAi plants, we found that not only the expression of *GmNIN1a*, but also the expression of other three *GmNINs,* was reduced, suggesting that our *GmNIN1a* RNAi plants phenocopy a *GmNIN* quadruple mutant. Moreover, knock down other *GmNINs* reduced nodule numbers, and gene expression showed these RNAi lines, also knock down other *GmNINs*. Our RNAi analysis of *GmNINs* suggest that they act in a functionally redundant manner to regulate the root nodule symbiosis. To further validate this hypothesis, we made a series of *GmNINs* mutants using a CRISPR-Cas9 approach. Unexpectedly, we found that *Gmnin1b* and *Gmnin1a nin1b* showed a wild type nodulation phenotype, including root hair deformation, rhizobial infection and nodule numbers. *Gmnin2a nin2b* mutant exhibited an intermediate phenotype with reduced nodule number. *Gmnin1a nin2a nin2b* triple and *Gmnin1a nin1b nin2a nin2b* quadruple mutants showed a typical *Ljnin* and *Mtnin* nodulation phenotype, including excessive root hair deformation, nearly no rhizobial infection events and no nodules. These results suggested that GmNIN1a, GmNIN2a, GmNIN2b are required for nodule organogenesis in a dosage dependent manner, while GmNIN1b is not able to compensate their function in *Gmnin1a nin2a nin2b* triple mutant.

Unexpectedly, *Gmnin2a nin2b* mutant showed a hyper infection phenotype that was similar to *Ljdephne* or *Mtdephne-like* mutant (Yono et al., 2014; Liu et al., 2019). Considering the non-infection phenotype in triple and quadruple mutants and normal infection in *Gmnin1a nin1b* double mutant, it is less likely that GmNIN2a and GmNIN2b simply play a negative role in rhizobial infection. One possibility is that the *GmNINs* expressions at cellular level might be differentiated with diversified biological implications. For instance, we speculate that *GmNIN2a* and *GmNIN2b* in nodule cortex cells might non-cell-autonomously modulate *GmNIN1a* expression in epidermal cells. In *Gmnin2a nin2b* mutant, *GmNIN1a* in epidermal cells might be induced and lead to more infection events. So far, we failed to clone *GmNIN1a* promoter for detailed expression analysis, and further studies are required to validate these hypotheses. Taken together, our results show that *GmNINs* play functionally redundant yet complicated roles in the root nodule symbiosis.

Due to multiple genome duplications, approximately 75% of soybean genes are duplicated leading to the perspective of high genetic redundancy (Schmutz et al., 2010). In some cases, duplicated genes are asymmetrically redundant, as mutation in one gene may cause a phenotype while in the other gene showed no or weak phenotype (Vaddepalli et al., 2019). Such asymmetric genetic redundancy is usually explained by diverged or unequal gene expression (Vaddepalli et al., 2019). Here we show that *GmNIN1b* plays a minor role in root symbiosis, even though its expression is comparable to the other three *GmNIN* homologs, but the most important 2^nd^ CE region was not conserved in *GmNIN1b*, implying its expression pattern may different with other NINs. Such divergence may also be attributed to variations in protein sequences, GmNIN1b was shorter than other GmNINs, and overexpression of *GmNIN1b* lead to marginal phenotypes. Elucidation of the protein sequences among *GmNIN* homologs may provide further sights into the evolution of NIN genes in legumes.

Consistent with *LjNIN*, overexpression of *GmNINs* reduced nodule numbers after rhizobia inoculation, and this suppression nodulation was systemic and dependent on AON pathway (Soyano et al., 2013; Soyano et al., 2014). However, *GmNIN1a* and its putative target genes (e.g. *GmRIC1*, *GmRIC2* and *GmNF-YA1*) were also induced in *GmNIN1a* overexpression plants. This result was different with *LjNIN* ectopic expression which down-regulated *NIN* expression and NIN activity (Soyano et al., 2014). This difference might have resulted from our use of the constitutive 35S promoter to drive *GmNIN1a* expression, rather than the DEX-inducible expression system used for the study of *LjNIN*. Moreover, we found that the overexpression of *GmNIN1a*, *GmNIN2a* or *GmNIN2b*, but not *GmNIN1b*, can form spontaneous malformed root nodule-like structures in the absence of rhizobia inoculation. We noted some cases where the structures that formed appeared to resemble hybrids between lateral roots and nodules, with proliferation of cortical cells around a central vascular bundle. The formation of nodules with central vascular bundles has been reported in several plant and rhizobial nodulation mutants (Imaizumi-Anraku et al., 2000; Guan et al., 2013), and better understanding of this phenomenon might give insights into how the peripheral vasculature particular to nodules forms. All these results support that GmNINs have similar important roles in root nodule symbiosis, and GmNIN1a, GmNIN2a and GmNIN2b function redundantly to regulate nodulation.

## MATERIAL AND METHODS

### Biological Materials and Growth Conditions

*Glycine max* Williams 82 and Huachun 6, rhizobial strain *B. japonicum* USDA110 was used in this study. *nark* mutant in Huachun 6 genetic background was previously generated by CRSIPR-Cas9 (Bai et al., 2020). For hairy root transformation, *Agrobacterium rhizogenes* strain K599 was used. *A. tumefaciens* strain EHA105 was used for expressing in *N. benthamiana*, and *A. tumefaciens s*train GV3101 was used for generating CRISPR-Cas9 stable mutants. Plasmids were transformed into *Escherichia coli* DH10B or DH5a for cloning.

### Analysis of Promoter of *GmNINs*

The sequence of *GmNINs’* promoters were extracted from Phytozome 12.1 (http://phytozome.jgi.doe.gov/pz/portal.html) and multiple alignment using website (http://www.ebi.ac.uk/Tools/msa/clustalo/) and the conserved regions and putative CYCLOPS binding site in *GmNINs’* promoters were searched in their promoter regions.

The *GmNIN2a* CE and CYC were amplified from Williams 82 leaf genomic DNA as template, the primer sequences are shown in Table S3. The PCR products and binary vector pK7FWG2-R which was modified by adding GUS gene before GFP, then using ClonExpress® multis One Step Cloning Kit (Vazyme), to generate p*GmNIN2a_CE-CYC_*:GUS, p*GmNIN2a_CYC_*:GUS and the ΔD1, D2 or D3 in p*GmNIN2a_CE-CYC_*:GUS. The constructs were confirmed by DNA sequencing and expressed in *A*. *rhizogenes* strain K599 for hairy root transformation. The transgenic roots were transferred into a vermiculite:perlite mixure (1:1) and inoculated with *B. japonicum* USDA110 5-7 days after transfer. The roots were harvested for GUS staining at indicated timepoints after inoculation. The stained nodules were sectioned and images were taken by light microscope (Nikon Eclipse Ni).

### Transient Expression of GFP-GmNINs in *N. benthamiana* Epidermal Cells

The *GmNINs* cDNA were amplified from *G. max* Williams 82 root cDNA using the primers shown in Table S2. The PCR products were cloned into pDONR207 and then recombined into the vector pK7WGF2 to generate *35S-GmNINs,* and then recombined this product into pUb-3xflag and p35S:mCherry (in order to replace its pUb promoter). The constructs were confirmed by DNA sequencing and transformed into *A. tumefaciens* strain EHA105, then co-infiltrated into *N. benthamiana* leaves plus p19. The images were taken by confocal microscopy with DAPI staining and protein expression levels were analyzed by western blot 60 hrs after infiltration.

### RNAi and Overexpression of *GmNINs*

For knock down *GmNINs* by RNAi strategy, specific fragments of *GmNINs* (about 150-200 bp) were amplified by PCR using *G. max* cDNA as template. The PCR products were cloned into pDONR207 and then recombined into pUb-GWS-GFP. For overexpression *GmNINs*, full-lengths *GmNINs* were amplified by PCR using *G. max* cDNA as template and cloned into pDONR207, then recombined into pK7WGF2. The primers used are showed in Table S3 and all the constructs in pDONR207 were sequenced. These constructs were introduced into *A*. *rhizogenes* K599 by electroporation and then expressed in *G. max* Williams 82 by hairy root transformation. The transgenic hairy roots were planted into a vermiculite:perlite (1:1) mixture and inoculated by *B. japonicum* USDA110 5-7 days after transfer. *GmNINs* overexpression hairy roots also were planted in a vermiculite:perlite (1:1) mixture without rhizobial inoculation. And the phenotypes were analyzed at the indicated time points.

### Generation of *Gmnin* CRISPR-Cas9 Mutants

To generate *Gmnin* mutants by CRISPR-Cas9, two small-guide RNAs targeting different regions of *GmNINs* were designed and cloned into the pGES201 vector. The pooled constructs were introduced into *A. tumefaciens* strain GV3101, which was then transformed into soybean cultivar Huachun 6 by *Agrobacterium*-mediated transformation as described (Bai et al., 2020). The mutant genotypes were identified using the Hi-TOM platform (Liu et al., 2019).

To observe *Gmnin* CRISPR-Cas9 infection phenotype, the mutants seedling were grown in a vermiculite:perlite mixture (1:1) and inoculated with *B. japonicum* USDA110 contains GUS reporter. The inoculated roots were harvested 7 days or 21 days after inoculation and the roots were stained by GUS (7 dpi), and phenotypes were observed under a light microscope (Nikon Eclipse Ni).

### Gene Expression Analysis

The *G. max* roots were harvested and total RNA was extracted using an RNAprep Plant plus Kit (Tiangen), and 1^st^ strand cDNA was synthesized using Transcript One-Step gDNA Removal and cDNA Synthesis super Mix Kit (TransGen Biotech.). RT-qPCR was performed on three biological replicates using an ABI StepOne PCR detection system with SYBR Green (Takara). *GmUbiquitin* (*SUBI-2.2*) expression was used as an internal control.

### Statistical Analysis

Statistical significance was analyzed by Student’s *t*-test (\**P* < 0.05, \*\**P* < 0.01) or one way ANOVA (non-parametric or mixed) and error bars indicate SE. Histograms were generated using GraphPad Prism 8.0 software.

## ACKNOWLEDGMENTS

We thank Prof. Min Wei (Lanzhou U. China) for providing USDA110 carrying GUS strain, and Prof. Jeremy Murray (CEMPS, China) for helpful editing on the manuscript. This work was supported by CAS grant (ZDRW-ZS-2019-2) and the National Key Research and Development Program of China (2016YFD0100702).

## CONFLICTS OF INTEREST

The authors declare that they have no conflicts of interest.

